# Feasibility of a surface-coated lung model for the quantification of active agent deposition for preclinical studies

**DOI:** 10.1101/639245

**Authors:** Philipp Dörner, Philipp M. Müller, Jana Reiter, Martin C. Gruhlke, Alan J. Slusarenko, Wolfgang Schröder, Michael Klaas

## Abstract

Multiple drug resistance (MDR) of a growing number of bacterial pathogens represents an increasing challenge in conventional curative treatments of infectious diseases. However, the development and testing of new antibiotics is associated with a high number of animal experiments. A symmetrical parametrized lung test rig allowing the exposure of air-passage surfaces to antibiotics was designed and tested to demonstrate proof-of-principle with aerosols containing allicin, which is an antimicrobial natural product from garlic. An artificial lung surface is coated with bacteria embedded in a hydrogel and growth inhibition is visualized by 3-(4,5-dimethylthiazol-2-yl)-2,5-diphenyltetrazolium bromide (MTT), that is reduced from colourless to the dark blue formazan in the presence of metabolically active, living cells. A nebulizer is used to generate the aerosols. The results show that allicin has an antibiotic effect as an aerosol and that the deposition pattern of the active agent occurred mainly around the carinal regions. The model represents an integral system for continuous, spatial detection of aerosol deposition and allows the analysis of bacterial behaviour and the toxicity of the active agent. In this way the deposition of antimicrobial aerosols on the bronchial surfaces is characterized in preliminary tests without any animal experiments.

## Introduction

Data from the U.S. Department of health and human services show that the number of deaths due to respiratory diseases has constantly increased in the USA over the last decades.^34^ In 2001 a crude death rate (deaths per 100,000 population) of 43.2 was reported, in 2010 a rate of 44.7, and in 2016 of 47.8, which accounted for approximately 5.6% of deaths from all causes. In 2016 51,537 patients died due to pneumonia or influenza, accounting for 1.9% of all deaths. In this context, antibiotic-resistant bacteria are a growing challenge in conventional curative treatments of infectious pulmonary diseases, making diseases such as pneumonia and tuberculosis increasingly serious and life-threatening. Hence, these diseases require new effective antibiotic treatments, including novel antibiotic classes and new application strategies to ameliorate the build-up of antibiotic resistance.

In general, antibiotics may be delivered to the target organ or tissue either orally by ingestion, intravenously by direct injection, or via the pulmonary route, i.e., inhalation. In the case of infectious lung diseases, the administration via the pulmonary route can be performed via metered-dose aerosols, nebulizers or powder inhalers, or potentially in the gas phase for volatile substances. The pulmonary route of administration of the active agent provides a number of advantages for lung diseases compared to oral or intravenous administration. For instance, it should be easier to achieve an effective dose at the site of action by direct inhalation, accompanied by a more rapid effect and a potential reduction of side effects as well as a lower administered dose. However, the approval procedure for new substances requires preclinical studies and animal experiments, both for toxicological tests and pharmacological studies. In preclinical studies, the long term toxic effects of repeated doses and acute toxicity after a single dose must be tested. Before animal tests for toxicity and clinical efficacy, pre-knowledge about the behaviour of drug-loaded aerosol particles in the lung flow and their deposition, as well as possible synergistic effects of combinations of antibiotics against the target pathogen, would be very helpful. Therefore, a symmetrical parametrized model of three lung bifurcations was designed in order to carry out such experiments.

The flow in the human lung is characterized by complex flow structures in an intricate geometry. Experimental investigations of the flow field in the human lung primarily include velocity measurements in the area between the trachea and the *Bronchi segmentales*, the tertiary bronchi. As shown by Geoghegan et al.,^14^ these investigations are often carried out *in vitro* by using transparent lung models, which are either reconstructed from two-dimensional or three-dimensional parameterized data sets, e.g., as in Weibel^32^ or Hammersley and Olson^16^, or are based on patient-specific CT data. Große et al.^15^ performed particle-image velocimetry (PIV) measurements under steady inspiration/expiration and under oscillatory flow for several Reynolds and Womersley numbers using a realistic lung model. It was shown that the flow in the first bifurcation between trachea and bronchia is highly three-dimensional. The authors stated that the critical Reynolds number, describing the onset of vortical structures in the bronchia, increases for higher Womersley numbers and lower Reynolds numbers. Soodt et al.^29^ focused on the second bifurcation and investigated the unsteady flow field in the same lung model using stereo scanning PIV. They showed that broader velocity profiles lead to an increased mass flux in the upper bronchus at higher Womersley numbers. Extensive investigations of the flow field in the first bifurcations of the tracheobronchial tree were conducted by Bauer et al.^7, 8^. Their PIV measurements in a Weibel/Horsfield-model showed the influence of the ventilation frequency on the unsteady three-dimensional flow structures. Jalal et al.^18^ and Banko et al.^5^ determined the three-dimensional velocity field in an idealized Weibel-lung model and in a realistic CT data-based geometry using magnetic resonance velocimetry (MRV). The latter measurements included the mouth and throat area and, thus, simulated realistic inflow conditions. The measurements focused on steady inspiration and time-averaged velocity fields and showed that idealized geometries tend to produce weaker secondary flows. However, all these investigations focused on the flow field itself and did not consider particle and aerosol transport and deposition.

Regarding particle transport and deposition in the lung and the absorption of active substances in the human respiratory tract, preclinical studies usually comprise experimental investigations based on different model classes. According to Nahar et al.^22^ and Fröhlich and Salar-Behzadi^13^, the models can be classified as *in vitro, in vivo*, and *ex vivo* models. *In vitro* models include lung models with different degrees of complexity and cascade impactors. The latter have been used for decades to make particle deposits qualitatively visible, as introduced by Yamada et al.^35^. Investigations with a dry and inflated swine lung were conducted by Morozov and Kanev^21^. For *in vivo* models, imaging techniques such as gamma scintigraphy, positron emission tomography (PET), single-photon emission computed tomography (SPECT), or magnetic resonance imaging (MRI) measure particle deposition directly in living animals using radionuclides or non-ionizing radiation, as shown by Conway^11^. Schittny et al.^27^ used phase contrast imaging to determine the deposition of 200 nm gold particles injected into the lungs of young adult rats by intratracheal instillation. The deposition of particles can also be analysed on the basis of tissue examinations, as reported by Yanamala et al.^36^

In summary, it can be stated that, to the best of our knowledge, there are no models available that simulate the flow field and the particle deposition in the respiratory tract of a realistic human lung geometry simultaneously. *In vitro* lung models allow the accurate replication of the geometry of the human lung for a detailed analysis of the flow field, but they are not suitable to investigate particle deposition coupled with lung physiology. In contrast, *in vivo* models allow an accurate analysis of particle deposition, but suffer from the disadvantage of species-specific reactions to particles. Moreover, the differences between the species with respect to lung physiology make it difficult to transfer the results of animal experiments to humans. Hence, new experimental methods are required to determine and quantify the local deposition of an active substance in the human lung, to analyse the medical indication of the substance, to quantify the deposition at the diseased position, and to determine the dose-dependent effectivity against a target microbe.

Infectious bacterial or fungal lung diseases often develop in humans with a weakened immune system (children, the elderly, chemotherapy-or HIV patients) and in patients with congenital immunodeficiency such as SCID (severe combined immunodeficiency, leading to chronic mucocutaneous candidiasis), COPD (chronic obstructive pulmonary disease), or cystic fibrosis. Those patients are susceptible to most common bacterial lung pathogens like *Streptococcus pneumoniae, Staphylococcus aureus, Pseudomonas aeruginosa, Acinetobacter baumannii*, and *Haemophilus influenzae*. Among the fungi associated with lung disease, *Aspergillus fumigatus* is one of the most common species. The aforementioned bacteria belong to the list published by the World Health Organization (WHO)^33^ with the most dangerous antibiotic-resistant bacteria. Small rodents such as mice, rats, and guinea pigs are used as models for the investigation of pulmonary infectious diseases. For mice, the lung or thigh is infected with the bacteria and the animals are treated by subcutaneous injection or oral uptake of the test substances. This procedure is one of the most realistic and application-oriented models for experimental treatments. Disadvantages are the pain experienced by the animals and the number required. Animal models are also used for testing the administration of drugs by inhalation, i.e., via the pulmonary route. The most commonly used are the *in vivo* mouse/rat inhalation models, in which the animals are treated in a chamber, as shown by Andes and Craig^2^, and the *ex vivo* rat/rabbit inhalation models, in which the isolated, perfused, and ventilated lung tissue (IPL) is treated, as described by Beck-Broichsitter et al.^9^ and Seeger et al.^28^. An alternative to the animal experiments are *in vitro* experiments with co-cultures of pathogenic bacteria and human or mammalian cells. The co-culture model is also suitable for the investigation of molecular mechanisms. The disadvantage is that the experiments are carried out in Petri dishes, i.e., the setup is quite different from the application *in vivo*.

The overview of the established models illustrates the urgent need for new animal-free and application-oriented models for infection experiments and inhalation treatments, since no established model is capable of realistically simulating the effects of drugs *in situ* on lung pathogens. Van der Poll and Opal^30^ show that pneumococcal pneumonia often begins in the upper airways, i.e., in the area of the main bronchus and the segmental bronchus. Thus, our model was constructed to recapitulate the 2^nd^, 3^rd^ and 4^th^ branches (bronchi) in the lung, in order to model deposition of active test substances in this critical region. Such information is essential for the development of new antibiotics, of which allicin represents a potential candidate. Allicin (diallylthiosulfinate) is a defence substance produced by garlic tissues when they are damaged. A garlic clove of approximately 10 g can produce up to 5 mg of allicin (Borlinghaus et al.^1^). Allicin has a broad spectrum of antimicrobial action and is effective against bacteria, fungi, oomycetes, and protozoa.^1, 3, 4, 12, 17, 24^ Allicin is effective against MDR strains of human lung pathogenic bacteria and because it is volatile it has an antibiotic effect in the gaseous phase (Reiter et al.^25^, Leontiev et al.^20^). Thus, allicin was chosen as the test substance for an inhalative aerosol in the lung model to show proof of principle for its function. To establish the potential for new antibiotic substances in the clinical situation prior to animal experiments, a detailed understanding of the fluid behaviour as well as the deposition behaviour of drug-loaded aerosol particles is required. The lung model reported here is a generic *in vitro* system to spatially analyse the particle deposition process and the effectiveness of the antibiotic substance with respect to a test bacterium. For ease of experimentation a non-pathogenic *E. coli* isolate was used, but given the appropriate laboratory safety category, there is no reason why lung pathogens could not be tested directly.

In this study, a first step towards a new method is described to test bacteria or human lung cells embedded in a hydrogel-coated generic lung model for preclinical active agent aerosol investigations. This allows the possibility of studying the antibiotic effectiveness as well as the pulmonary toxicity, if in a further development cultured lung epithelial cells can be used to line the model. For this purpose, vitality and proliferation and, if necessary, other physiological parameters can be tracked in real time using genetically coded reporter constructs. This is not easily possible in animal models. With the data, generated under realistic *in vitro* conditions, detailed treatment routines can be conceived before progressing to animal experiments. This would reduce the number of animal tests needed.

## Materials and Methods

### Statistical lung model

For a comprehensive analysis of the active agent deposition, the flow measurement methods, i.e., the particle-image velocimetry (PIV), and the biological measurements for the lung model are to be described. This study focuses on the feasibility concerning the biological method of wall-embedded bacteria and reporter and detection systems. The flow methods are also considered by using the same model characteristics for PIV-capable transparent models to measure the deposition.

### Material test for allicin resistance

Allicin was synthesized as previously described by the oxidation of diallyl-disulfide with hydrogen peroxide^1^. The materials which were considered for the construction of a lung model were tested for their chemical resistance against allicin. For this purpose, 20 μl 96 mM allicin was applied to the material surface, covered by the lid of a Petri dish, and incubated at 37 °C for 18 h. After incubation, the surface was cleaned with water and viewed under the microscope. Allicin-resistant materials are stainless steel (SST), soda lime glass (SLG), polyamide (PA), polyethylene terephthalate (PET), polyoxymethylene (POM), and polytetrafluorethylene (PTFE). The surfaces of polystyrene (PS) and polyvinyl chloride (PVC) showed a pitting reaction to allicin and are therefore chemically unstable. Polymethyl methacrylate (PMMA), polyethylene (PE), and polylactic acid (PLA) became opaque, whereas aluminium (Al) became darkly discoloured. The surface of polyether ether ketone (PEEK) became dark and rough (Table 1). Due to the chemical resistance and the good machining properties, POM was chosen as the model material.

**TABLE 1.** Chemical resistance of different materials to 96 mM allicin applied on the surface for 18 h at 37 °C. The symbols +, -define no impact of allicin on the surface and surface destruction, respectively.

| material | resistance | material | resistance | material | resistance |
| --- | --- | --- | --- | --- | --- |
| Al | - | PA | + | PLA | - |
| SST | + | PE | - | PS | - |
| SLG | + | PEEK | - | PVC | - |
| PMMA | - | PET | + | POM | + |
|  |  |  |  | PTFE | + |

### Model geometry

For a first detailed analysis of the active agent deposition, the complexity was reduced by neglecting the complex inflow conditions of the larynx. Instead, a fully developed laminar and steady pipe flow is used. The location of pneumococci is generally in the upper airways, i.e., in a range of the second to the tenth generation bronchi. For increasing generations, the bronchial diameter decreases exponentially. Geometrical scaling, see Figure 1, is possible by considering the similarity numbers of the flow and the aerosol. This leads to scaled particles, since the Stokes number (St)

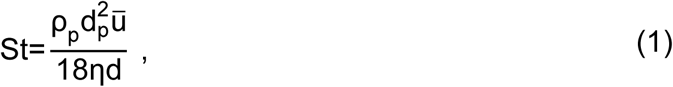

determined by the particle density ρ_p_, the particle diameter d_p_, the mean flow velocity 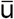, the dynamic viscosity η, and the diameter of the pipe d, is proportional to the scaling factor. The same applies for, the Reynolds number (Re)

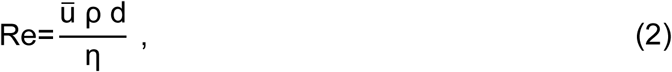

where ρ represents the density of the fluid. The Dean number (De), defined as

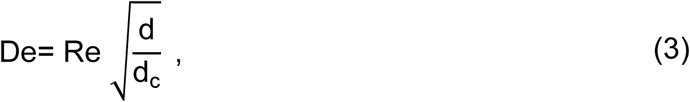

with d_c_ as the diameter of the curvature, is a measure for secondary flow pattern in curved vessels with a small curvature ratio. However, with respect to the investigation of drug delivery with wall embedded bacteria, a larger particle size would result in a change in the bacterial killing rate, since the bacteria cannot be scaled simultaneously. Thus, in this study a scaling is avoided and the second to the fifth generation at life-size were chosen.

**FIGURE 1.**
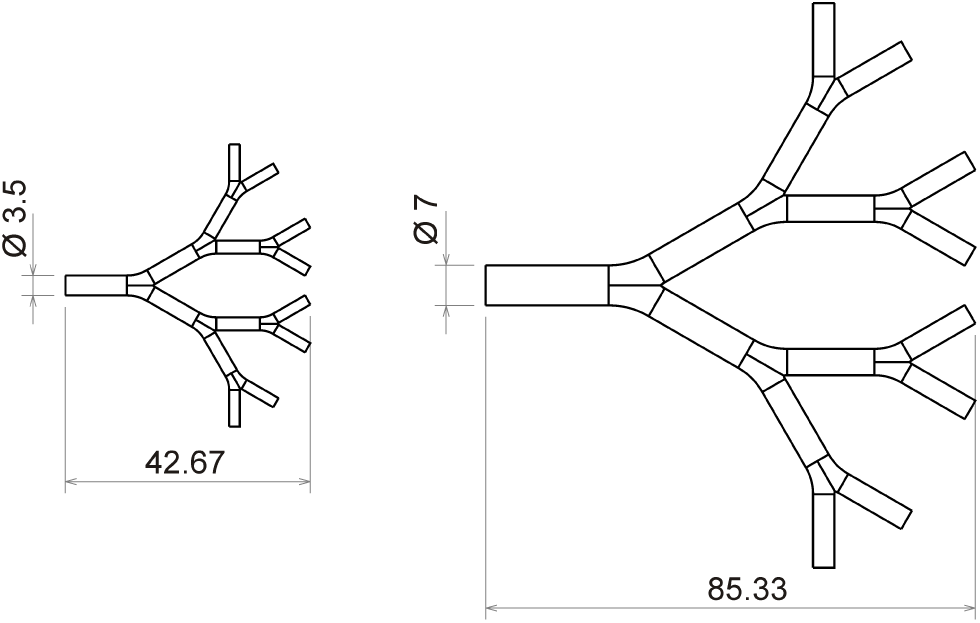
Comparison of the original size in [mm] of the 5^th^ – 8^th^ generation (left) and a 2:1 scaled model (right).

The geometric parameters of the lung model are defined in Figure 2. They are adapted from physiological values of Hammersley and Olson^16^ and Weibel^32^.

**FIGURE 2.**
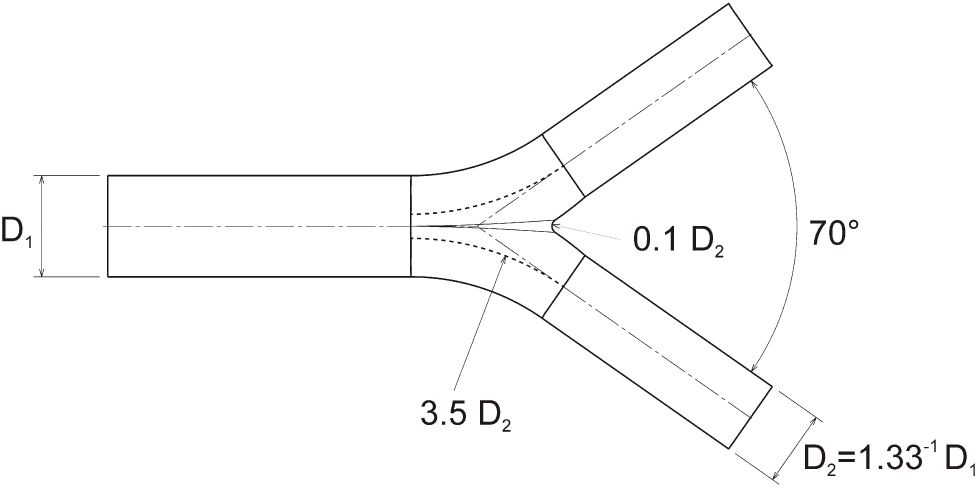
Parameterized symmetric lung model geometry from Hammersley and Olson^16^ with L/D_1_=3.

The daughter vessel diameters D_2_ are scaled by the mother vessel diameter D_1_ with a factor of 1.33^-1^. Furthermore, the daughter branch radius is parameterized by 3.5 D_2_. The carinal flow divider radius is scaled by 0.1 D_2_, the length by L/D_1_=3, and the angle between the daughter vessels is constant at 35°. Table 2 shows the parameters of the lung model with the resulting Re, St, and De numbers.

**TABLE 2.**
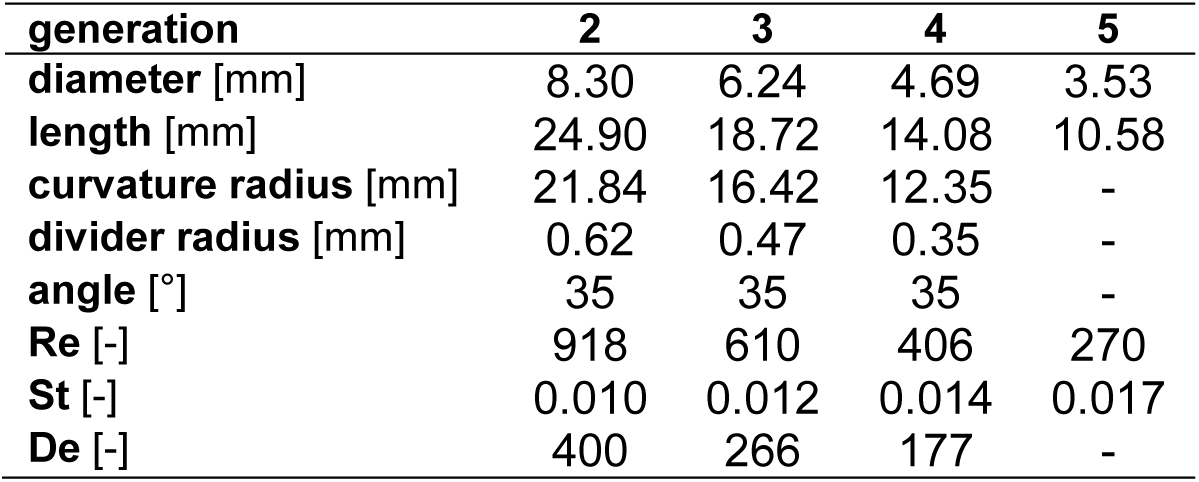
Dimensions and parameter of the parameterized lung model.

Figure 3 (left) depicts the Reynolds and Stokes numbers for Weibel’s model for the first 15 generations. Based on this distribution, the Reynolds and Stokes numbers for the parameterized model of the current study are defined. Since the daughter vessels are scaled by the Hammersley and Olson^16^ parameters, Stokes and Reynolds number differ from the Weibel model. Note that Hammersley and Olson^16^ also used anatomic data from four human lungs to developed a further detailed approximation of smaller airways. The anatomy of the lungs depends on various factors, e.g., sex, age, and medical condition.

**FIGURE 3.**
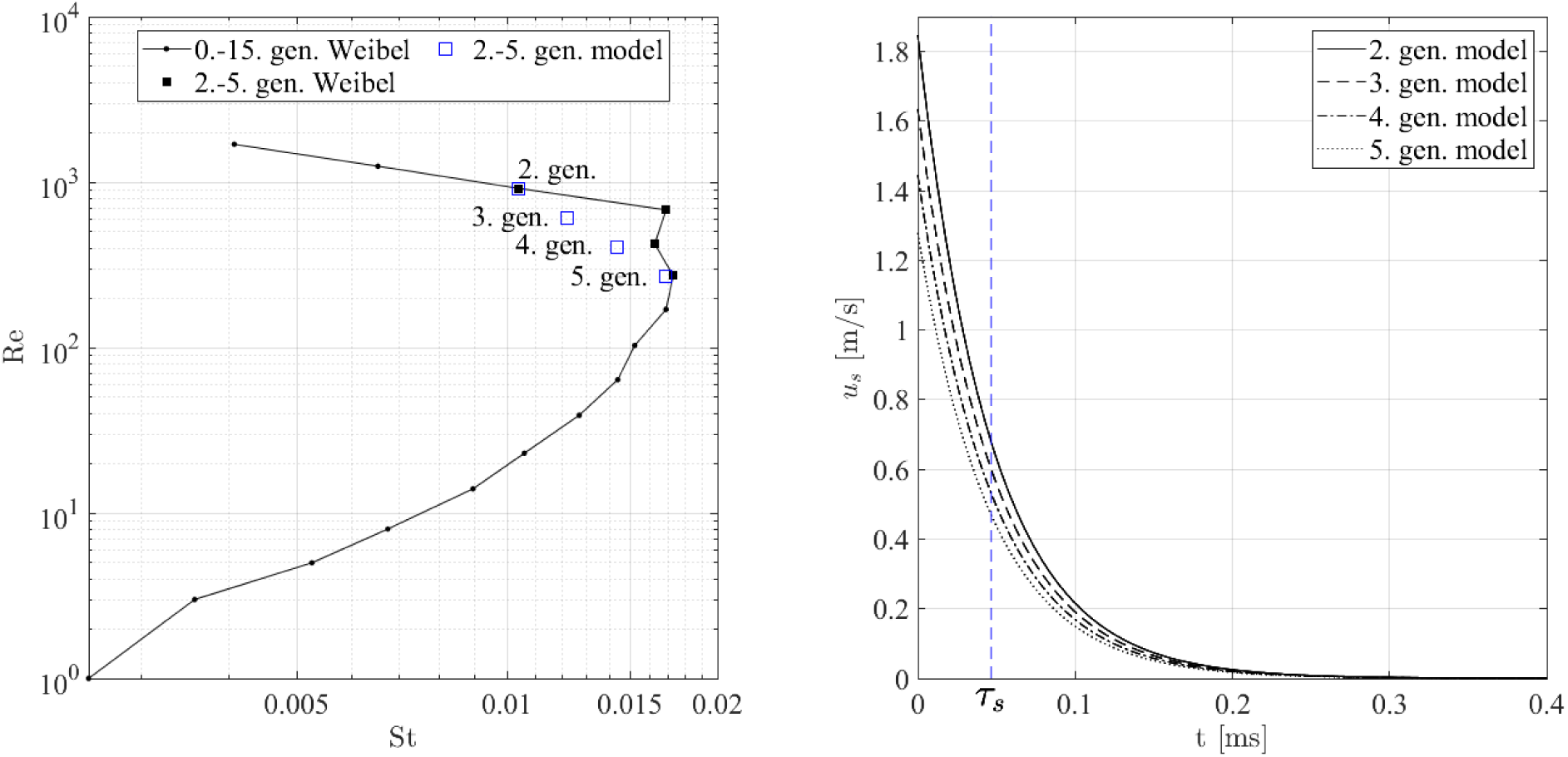
Reynolds and Stokes numbers for the Weibel model (black) and the parameterized lung model (blue) for a particle diameter of 4 μm, a volume flow of 24 l min^-1^, air at 37 °C, particle medium as water at 20 °C, and ambient pressure of 101 325 Pa (left). The slip velocity u_s_ and the relaxation time τ _s_ under decelerating conditions (right).

Zhang et al.^37^ simulated a two-phase lung flow, i.e., particle and air, from the third to the sixth lung generation of a Weibel adapted lung model under different breathing rates and compared their results to the experiments of Kim and Fisher^19^. It was shown that the deposition efficiency increases strongly with higher Stokes number. The measurements in this study are performed in this parameter range such that particle deposition can be expected. Figure 3 (right) shows the particle response time

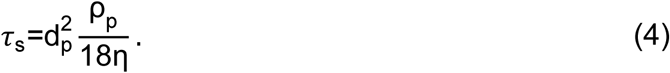

for a particle with a diameter of d_p_= 4 μm as they are used here. The slip velocity u_s_ of a decelerating flow is defined as

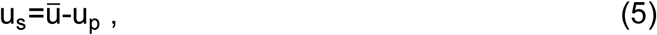

where u_p_ represents the particle velocity

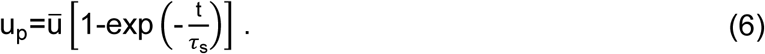

It is shown that the relaxation time *τ*_s_, i.e., 63% of 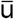, is reached at 0.0466 ms and is within the order of magnitude of Zhang et al.^37^.

### Surface coating

To spatially and continuously detect the aerosol deposition for the analysis of the bacteria behaviour and the toxicity, i.e., the concentration of the active agent, and to develop additional biological reporter and detection systems, a surface coating with embedded bacteria might be a solution with several biological as well as fluid mechanical benefits. With the local independence of the biological detection system within the lung model, bacteria growth and aerosol effects can be measured by extraction of a small sample.

For the surface coating the shape stability, the durability, i.e., preventing drying-out, the demouldability, and the surface quality must be ensured for a culture medium with bacteria. Preliminary tests with a curved layer of 1 mm thickness of agar (Agar-Agar Kobe I, powdered, for microbiology, Carl Roth, Germany) and agarose (UltraPure™ Agarose, Invitrogen™) showed that either an agar concentration of 1.5% or an agarose concentration of 0.7% satisfied these requirements. To dissolve the powders into water the mixtures were heated up in a microwave to 85°C and cooled down to 50°C before casting. Non-pathogenic *E. coli* bacteria were used as a test system embedded into the agar coating of the lung model. For culturing, a LB (Luria–Bertani) culture medium from Bertani^10^ with 1.5% (w/v) of agar was used. The ingredients were mixed with distilled water and autoclaved for 20 min at 121°C. Subsequently, the molten LB medium was tempered in a water bath at 50°C and the bacteria were added before casting.

In Figure 4, the casting process of the surface coating is shown. Initially, a negative mould has to be manufactured to cast a stamp with the inner, final curvature of the bifurcation system with an offset of 1mm. The negative mould was milled from aluminium and has a raised step contour parallel to the geometry. These steps result in a groove directly next to the curvature of the stamp and prevent shrinkage effects of the agar layer and leakage. Furthermore, the outlets of the mould are broadened due to the direct contact to the environment. The stamp was cast from epoxy resin L with a hardener W 300 (R&G Faserverbundwerkstoffe GmbH, Germany). Before casting the agar surface, the POM moulds and the connection elements are sterilized by UV light for 20 min. In the next step, the tempered agar-bacteria mixture (20 ml LB-agar and 300 μl bacterial suspension (optical density by wavelength 600 nm, OD_600_= 0.2) is quickly filled into the mould. Immediately afterwards, the stamp is pushed down by the precise positioning of dowel pins. The stamp is held for 5 min to ensure complete agar solidification. Next, the stamp is removed and excess medium at the inlet and outlets are cut off with a scalpel to obtain sharp inlet and outlet contours. This procedure is repeated for the other half mould before the upper and lower halves are assembled.

**FIGURE 4.**
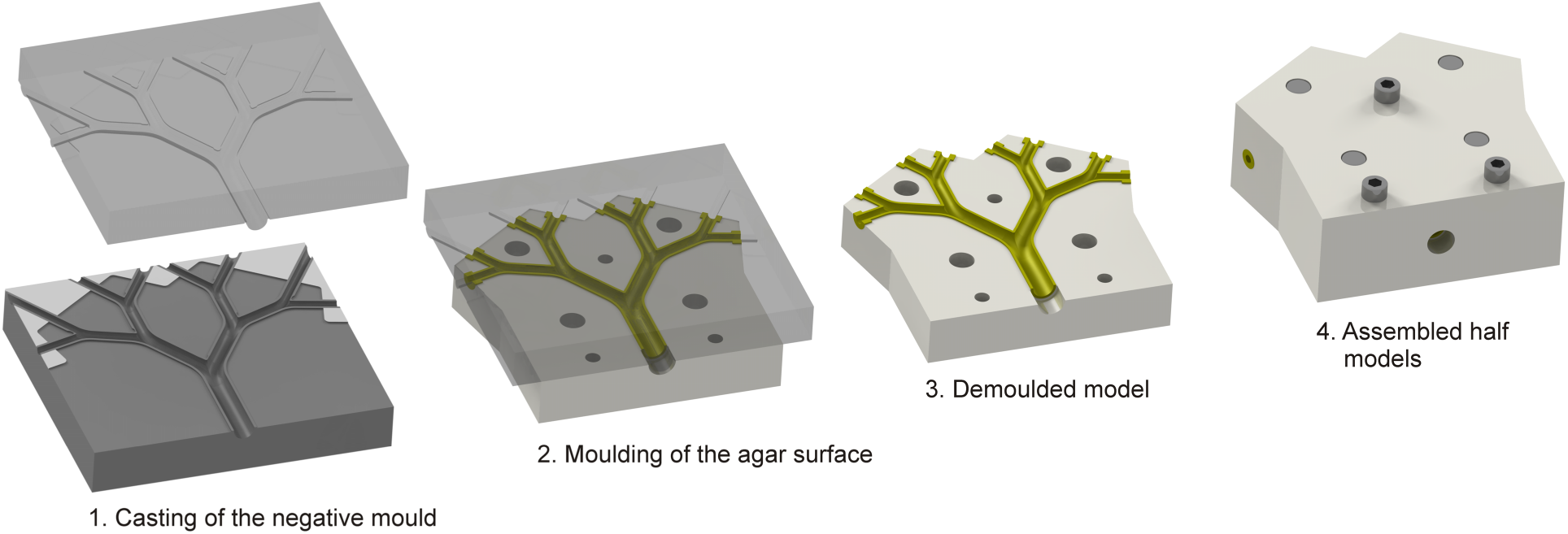
Moulds and casting process of the agar coated lung model.

### Experimental Setup

Figure 5 shows a sketch of the experimental setup. A steady flow is generated by an aerosol generator and air-supply. To ensure well defined inflow conditions, a laminar inlet tube made of quartz glass with an inner diameter of 8.3 mm and a length to diameter ratio of 55 is mounted between the nebulizer and the lung model. The model is placed in a rectangular container which is ventilated and temperature controlled at 37°C to approximate conditions in the human body. The inlet tube is encapsulated by a PMMA tube which is connected to the container and equilibrated to 37°C. At the beginning of each experiment the flow rate is adjusted by a flow meter (Flo-Rite Model MR3000, Key Instruments) to 6 l min^-1^. The total flow rate, i.e., the flow generated by the aerosol generator and the virtually patient inspiration flow substituted by the supply air, is adjusted before the measurements, since the aerosol particles would deposit in the flow meter. The exit ventilation port on the incubator was fitted with a 0.45 µm filter. The whole setup is placed under a laboratory fume hood.

**FIGURE 5.**
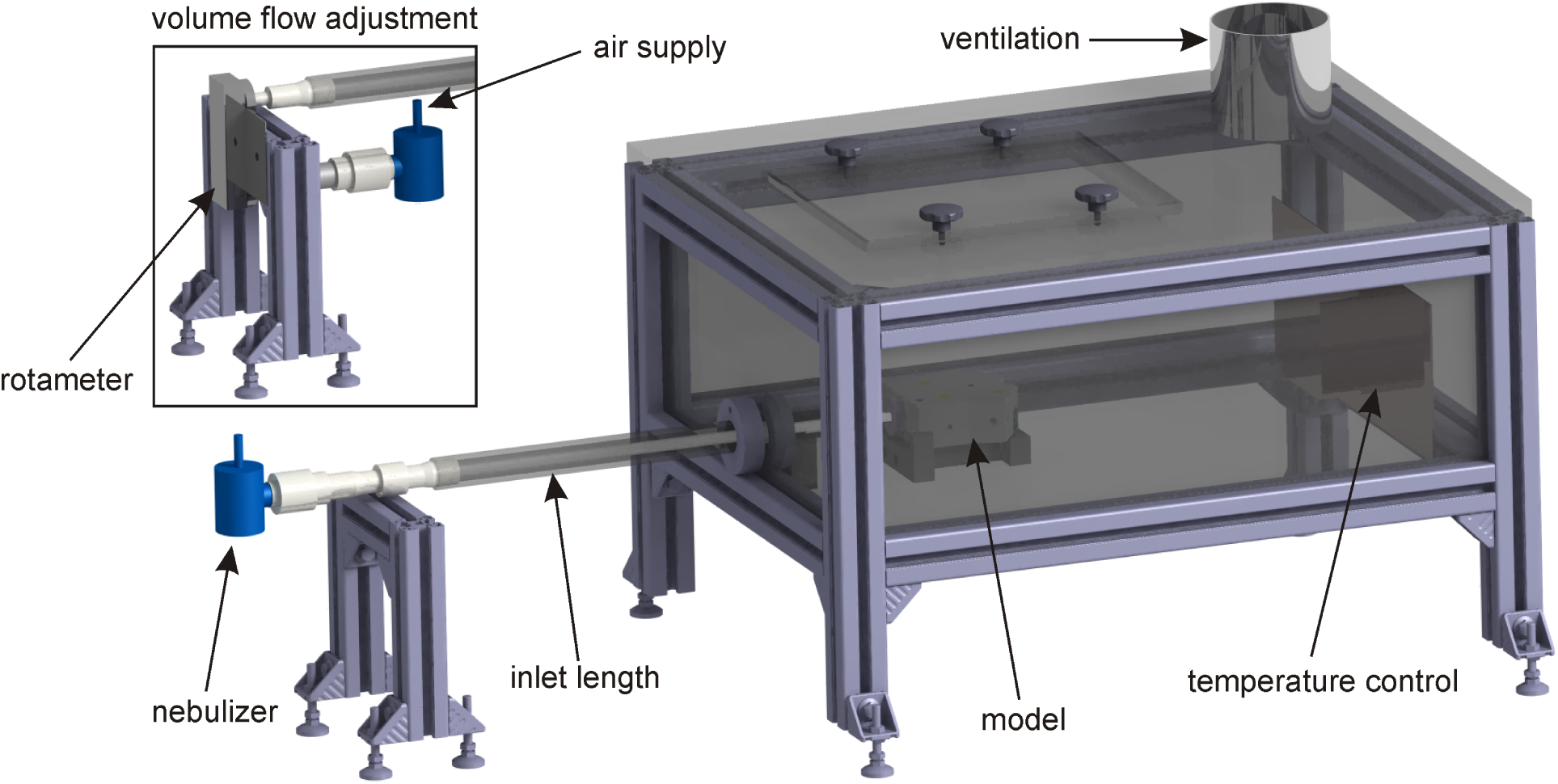
Experimental setup.

### Active agent aerosol generation

To ensure a particle generation that resembles medical applications, the PARI TurboBOY SX (PARI GmbH, Germany) aerosol generator is used with the PARI LC SPRINT nebuliser. This nebulizer possesses a patented peak inspiratory flow-control (PIF-control), which prevents high inspiratory velocities with inefficient particle deposition. For steady inflow conditions, the PIF-control module is removed and replaced by a sealed supply air connection where the volume flow can be adjusted by a pressure regulator. For this device, a mass median aerodynamic diameter (MMAD) of ≈ 4 μm is reached at 6 l min^-1^. Wang et al.^31^ measured a MMAD of 4.068 μm at 20 l min^-1^.

## Results

First, the validity of the test system using MTT to locate metabolically active bacteria was tested. To do this the agar-embedded bacteria were exposed to a water aerosol. The treatment was performed within the measuring container which was preheated at 37°C. After the casting process and the flow rate adjustment to 6 l min^-1^, the model was connected to the inlet tube and placed in the container. The nebulizer was filled with 8 ml water. The test was stopped after 20 min, when the water was entirely nebulized. After treatment the lung model was covered with a moist paper towel and aluminium foil to prevent drying-out and was incubated for 18 h at 37°C. Afterwards, the model halves were separated and the agar layer was sprayed with a 0.5% MTT solution to test for metabolic activity. The model halves were covered with PE foil and incubated at 37°C for 30 min. The MTT solution was reduced by the living cells giving a dark blue-purple colouration due to the production of insoluble formazan. After treatment with a water aerosol the whole hydrogel showed a dark blue colouration, demonstrating that living bacteria were uniformly present in all parts of the model (Figure 6 left, centre).

**FIGURE 6.**
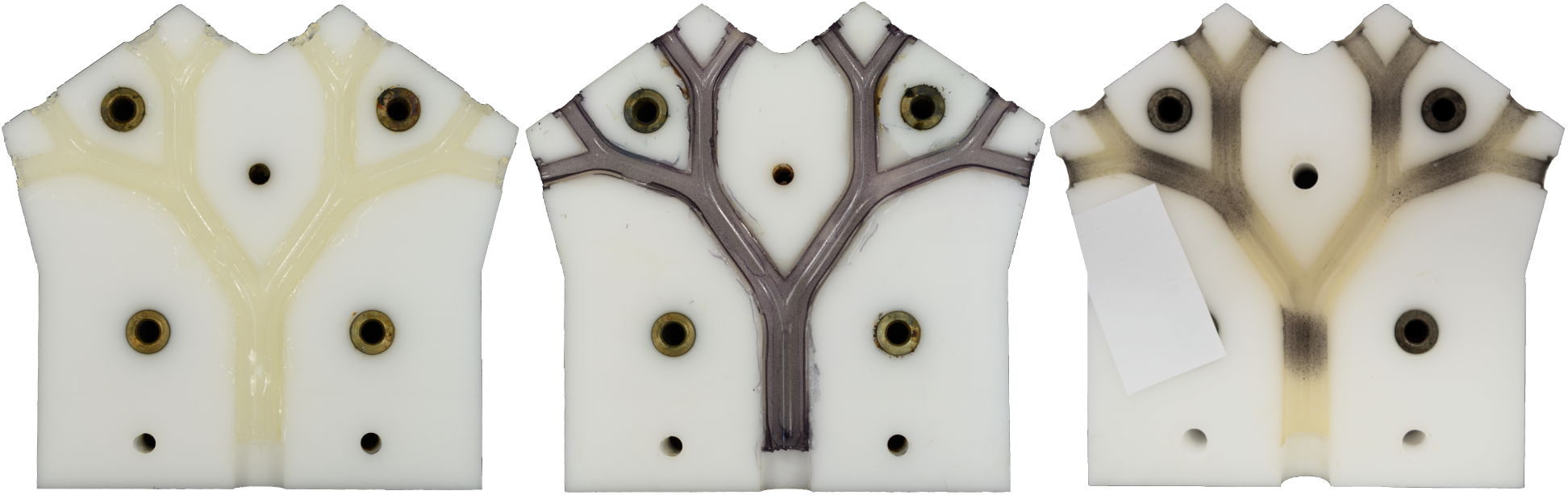
Controls: The lung model with a surface coating of bacteria-seeded agar was treated for 20 min with a nebulized water aerosol and incubated at 37°C for 18 h. The appearance before (left), and after (centre) spraying with tetrazolium salt MTT is shown. The dark colouration shows the even distribution of living bacterial cells throughout the model. Inhibition of bacteria after treatment with a 0.6 mM allicin aerosol for 20 min can be seen as pale non-stained regions (right).

Secondly, the deposition pattern of an antibacterial test substance as it would occur in the secondary, tertiary and quaternary bronchi is shown in Figure 6 (right) using an allicin aerosol. The regions of inhibition can be clearly seen as pale areas after spraying with MTT.

Next the model was used to show dose-dependency for the effectivity of an antibacterial agent using treatments with 0.5 mM, 0.6 mM, and 0.75 mM allicin aerosols. Figure 7 shows the deposition and inhibition of bacteria of the upper half of the model and demonstrates a clear dose-dependency of inhibition for the three allicin concentrations. The greyscale images in the centre plane in Figure 7 are normalized by their maximum intensity value. The top and bottom figures show fused images of allicin concentration results of 0.5 mM/0.6 mM (bottom left), 0.6 mM/0.75 mM (bottom right), and 0.5 mM/0.75 mM (top). To show the antibiotic effect in relation to the concentration level, the RGB composite images represent two overlaid normalized greyscale images of the selected concentration combination. They show the difference in varying colour scaling. This is achieved by placing one image into the green colour channel and the superimposed image into the red and blue channel. The same intensity level of both images is characterized by grey areas, whereas magenta and green show regions where the image intensities differ.

**FIGURE 7.**
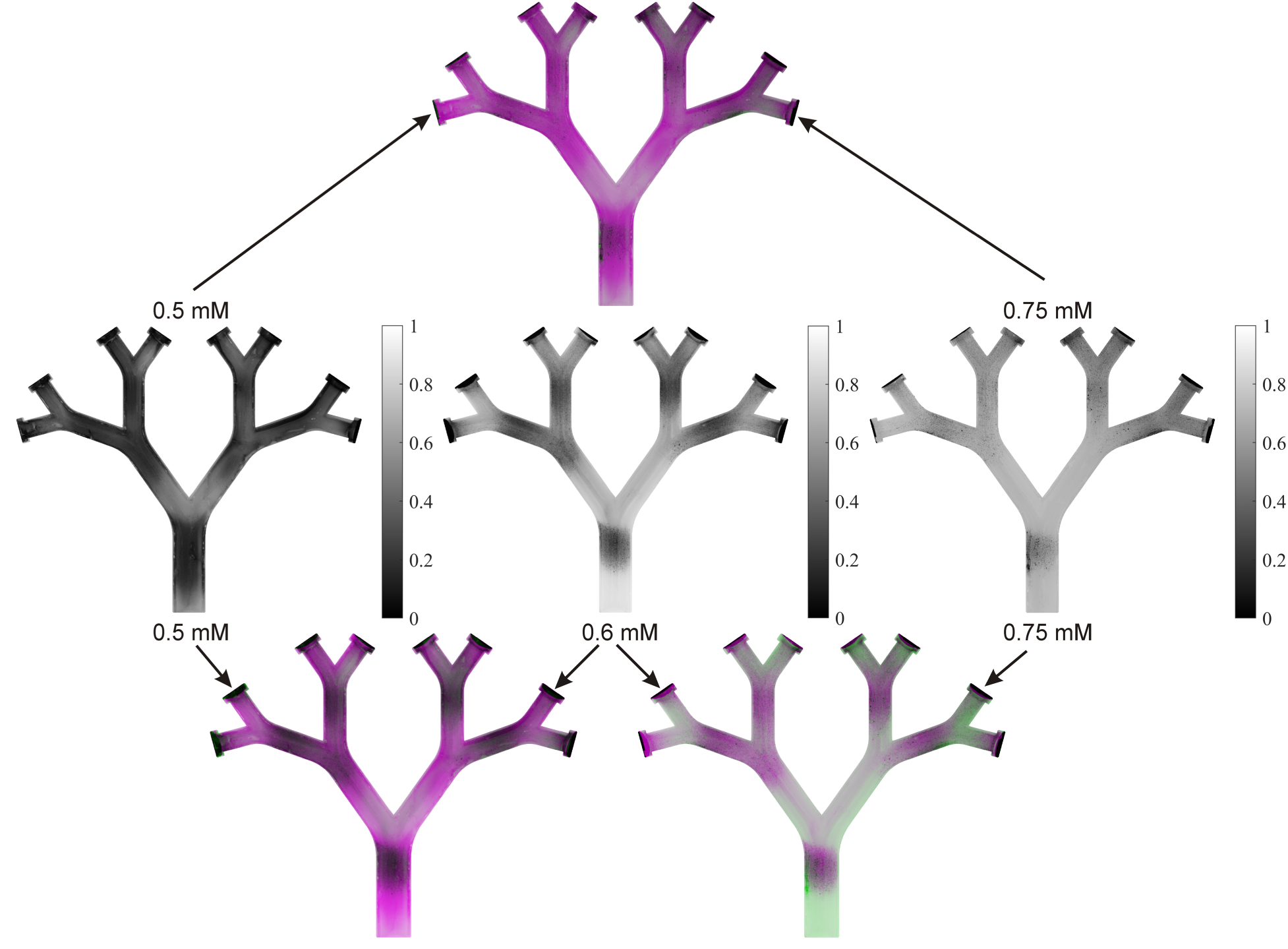
Deposition pattern of aerosols of 0.5 mM, 0.6 mM, and 0.75 mM allicin represented by normalized greyscale images (centre). The centre row shows a clear dose-dependency for bacterial inhibition by the allicin aerosol seen as decreasing area of formazan production from left to right. Composite images (top, bottom) overlaid by different colour scales, where grey regions indicate the same intensity of both images, magenta and green represent regions where the intensities are different.

## Discussion

Since individuals have different characteristic lung anatomy, the inhalation of active agent aerosols will be patient-specific. That is, it depends on many factors such as the medical condition, the clinical conditions, the nebulizer design, variations in the breathing and the tidal volume which is why the peak inspiratory flow rate has to cover a broad range. A flow rate of 6 l min^-1^ at the inlet of the second lung generation (first model generation) was chosen, which corresponds to 24 l min^-1^ inspiratory peak flow. This is within the range of the measured breathing pattern of Roth et al.^26^, Nikander et al.^23^, and Bauer et al.^6^

Figure 7 (centre) evidences that the treatment with allicin as an aerosol shows a reduction of the bacterial vitality for all tested concentrations. It is clearly visible that the test solutions inhibit the bacteria in a dose-dependent manner. According to the results of Zhang et al.^37^, the particle deposition occurs mainly around the carinal regions, where the flow bifurcates into the daughter branches. Slight asymmetries due to a slight misalignment of the model with respect to the inlet tube are reflected in the deposition pattern of the right half of the model being slightly more pronounced for 0.6 mM and 0.75 mM allicin than in the left half. Nevertheless, the deposition is more or less similar in both halves. The transition region from the inlet tube to the model exhibits small areas of non-living bacteria. This could be caused by small imperfections of the inlet-model transition. Despite low St numbers and suitable horizontal positioning of the model, the deposition pattern of the bottom half exhibits larger areas of dead bacteria than the upper half of the model. This could be caused by the gravitational force on the aerosol causing higher deposition rates leading to increased killing rates of the bacteria in the lower half. Neglecting the slight asymmetry at the model-inlet tube, the first bifurcation of the model shows a symmetrical deposition pattern for all concentrations of allicin tested. Due to the decomposition of the symmetrical velocity profile from the mother vessel and the curvature of the bifurcation, a secondary flow occurs. This is evident by the asymmetrical deposition in the daughter vessels in Figure 7 (centre) at 0.5 mM allicin, where the inhibition is much stronger at the inner wall of the bifurcation. The slight misalignment in Figure 7 (centre) at 0.6 mM yields the higher velocity at the inner wall, which contributes to a longer deposition length in the right daughter vessel of the first model generation. The skewed velocity profile in the first and second bifurcation influences the symmetry of the inner third model bifurcations as shown at 0.5 mM in Figure 7 (centre).

It can be summarized that allicin has showed a dose-dependent antibiotic effect as an aerosol and its distribution in the lung model realistically models that which might be expected by inhalation by a test animal. This is shown in the image comparison in Figure 7 (top, bottom). The comparison of 0.5 mM/0.6 mM exhibits a higher difference in dead area compared to 0.6 mM/0.75 mM as shown by the magenta colouration, where the deposition area at the carinal regions for the 0.5 mM/0.6 mM is extended. The comparison of 0.6 mM and 0.75 mM results demonstrates that the pattern of inhibition at the first bifurcation is almost identical for both concentrations. Differences occur mainly in the second and third bifurcations. Due to an almost analogous intensity distribution the differences are smaller compared to the lower concentration, which results in less saturation of magenta and the occurrence of green. The comparison of the highest and lowest allicin concentration is shown in Figure 7 (top). It is clearly visible that the deposition area, of inhibited, i.e., the dead bacteria, is much more pronounced for the highest concentration of test substance (i.e. allicin).

It should be mentioned that these steady flow results within a symmetrical lung model do not of course represent all the extra complexities of real human lung flow, with surface irregularities and the presence of bacterial conglomerates. However, the results illustrate a qualitative prediction for the aerosol deposition behaviour in bifurcated flows and represent a first step towards a new method for preclinical testing of an active agent. Furthermore, by using a transparent comparative model for PIV applications with refractive-index-matched fluids, the flow behaviour can be investigated in detail simultaneously.

## Conclusion

The feasibility of a bacteria-embedded hydrogel coated generic lung model for preclinical testing of active agent aerosol deposition was investigated. The literature shows that allicin has antibiotic effects and it was used as an aerosol to establish a model to determine the efficiency and quality of the approach. A symmetrical statistical lung model was designed taking into consideration the non-scalability of the embedded test bacteria and the chemical resistance of the material. A casting process of the hydrogel surface coating is described and the embedding of living bacteria was confirmed. It was shown that allicin as an aerosol has an effective impact on the cells and the deposition of the active agent for different concentrations is described. This study is the first step towards a new method of preclinical active agent aerosol investigations. Using this model avoids animal sacrifice for preliminary testing of new antibiotics to combat lung-pathogens by inhalation. In the future, the complexity of the inflow conditions, the three dimensionality of the curvature, the effects of evaporation, the embedding of biological reporter detection systems, and the diffusion process are to be investigated. Using this model will enable different sensitivities of test bacteria to be visualized and the synergistic effects of antibiotic combinations to be tested before animal experiments are required. This will lead to a reduction of pre-clinical animal testing.

## References

1. Albrecht F., R. Leontiev, C. Jacob and A. J. Slusarenko. An optimized facile procedure to synthesize and purify allicin. Molecules 22:770, 2017.

2. Andes D. and W. A. Craig. Pharmacodynamics of the New Fluoroquinolone Gatifloxacin in Murine Thigh and Lung Infection Models. Antimicrob. Agents Chemother. 46:1665–1670, 2002.

3. Ankri S. and D. Mirelman. Antimicrobial properties of allicin from garlic. Microb. Infect. 1:125–129, 1999.

4. Arora D. S. and J. Kaur. Antimicrobial activity of spices. Int. J. Antimicrob. Agents 12:257–262, 1999.

5. Banko A. J., F. Coletti, D. Schiavazzi, C. J. Elkins and J. K. Eaton. Three-dimensional inspiratory flow in the upper and central human airways. Exp. Fluids 56:117, 2015.

6. Bauer A., P. McGlynn, L. L. Bovet, P. L. Mims, L. A. Curry and J. P. Hanrahan. The Influence of Breathing Pattern During Nebulization on the Delivery of Arformoterol Using a Breath Simulator. Respir. Care 54:1488–1492, 2009.

7. Bauer K. and C. Brücker. The Influence of Airway Tree Geometry and Ventilation Frequency on Airflow Distribution. J. Biomech. Eng. 137:081001, 2015.

8. Bauer K., A. Rudert and C. Brücker. Three-Dimensional Flow Patterns in the Upper Human Airways. J. Biomech. Eng. 134:071006, 2012.

9. Beck-Broichsitter M., J. Gauss, C. B. Packhaeuser, K. Lahnstein, T. Schmehl, W. Seeger, T. Kissel and T. Gessler. Pulmonary drug delivery with aerosolizable nanoparticles in an ex vivo lung model. Int. J. Pharm. 367:169–178, 2009.

10. Bertani G. Studies on lysogenesis. I. The mode of phage liberation by lysogenic Escherichia coli. J. Bacteriol. 62:293–300, 1951.

11. Conway J. Lung imaging - Two dimensional gamma scintigraphy, SPECT, CT and PET. Adv. Drug Del. Rev. 64:357–368, 2012.

12. Curtis H., U. Noll, J. Störmann and A. J. Slusarenko. Broad-spectrum activity of the volatile phytoanticipin allicin in extracts of garlic (Allium sativum L.) against plant pathogenic bacteria, fungi and Oomycetes. Physiol. Mol. Plant Pathol. 65:79–89, 2004.

13. Fröhlich E. and S. Salar-Behzadi. Toxicological Assessment of Inhaled Nanoparticles: Role of in Vivo, ex Vivo, in Vitro, and in Silico Studies. Int. J. Mol. Sci. 15:4795–4822, 2014.

14. Geoghegan P. H., N. A. Buchmann, C. J. T. Spence, S. Moore and M. Jermy. Fabrication of rigid and flexible refractive-index-matched flow phantoms for flow visualisation and optical flow measurements. Exp. Fluids 52:1331–1347, 2012.

15. Große S., W. Schröder, M. Klaas, A. Klöckner and J. Roggenkamp. Time resolved analysis of steady and oscillating flow in the upper human airways. Exp. Fluids 42:955–970, 2007.

16. Hammersley J. R. and D. E. Olson. Physical models of the smaller pulmonary airways. J. Appl. Physiol. 72:2402–2414, 1992.

17. Ilić D. P., V. D. Nikolić, L. B. Nikolić, M. Z. Stanković, L. P. Stanojević and M. D. Cakić. Allicin and related compounds: Biosynthesis, synthesis and pharmacological activity. Facta Universitatis Series: Physics, Chemistry and Technology 9:9–20, 2011.

18. Jalal S., A. Nemes, T. Van de Moortele, S. Schmitter and F. Coletti. Three-dimensional inspiratory flow in a double bifurcation airway model. Exp. Fluids 57:148, 2016.

19. Kim C. S. and D. M. Fisher. Deposition Characteristics of Aerosol Particles in Sequentially Bifurcating Airway Models. Aerosol Sci. Technol. 31:198–220, 1999.

20. Leontiev R., N. Hohaus, C. Jacob, M. C. H. Gruhlke and A. J. Slusarenko. A Comparison of the Antibacterial and Antifungal Activities of Thiosulfinate Analogues of Allicin. Sci. Rep. 8:2018.

21. Morozov V. N. and I. L. Kanev. Dry Lung as a Physical Model in Studies of Aerosol Deposition. Lung 193:799–804, 2015.

22. Nahar K., N. Gupta, R. Gauvin, S. Absar, B. Patel, V. Gupta, A. Khademhosseini and F. Ahsan. In vitro, in vivo and ex vivo models for studying particle deposition and drug absorption of inhaled pharmaceuticals. Eur. J. Pharm. Sci. 49:805–818, 2013.

23. Nikander K., J. Denyer, N. Smith and P. Wollmer. Breathing Patterns and Aerosol Delivery: Impact of Regular Human Patterns, and Sine and Square Waveforms on Rate of Delivery. J. Aerosol Med. 14:327–333, 2001.

24. Rabinkov A., T. Miron, L. Konstantinovski, M. Wilchek, D. Mirelman and L. Weiner. The mode of action of allicin: trapping of radicals and interaction with thiol containing proteins. Biochimica et Biophysica Acta (BBA) - General Subjects 1379:233–244, 1998.

25. Reiter J., N. Levina, M. van der Linden, M. Gruhlke, C. Martin and A. J. Slusarenko. Diallylthiosulfinate (Allicin), a Volatile Antimicrobial from Garlic (*Allium sativum*), Kills Human Lung Pathogenic Bacteria, Including MDR Strains, as a Vapor. Molecules 22:2017.

26. Roth A. P., C. F. Lange and W. H. Finlay. The Effect of Breathing Pattern on Nebulizer Drug Delivery. J. Aerosol Med. 16:325–339, 2003.

27. Schittny J. C., S. F. Barré, R. Mokso, D. Haberthür, M. Semmler-Behnke, W. G. Kreyling, A. Tsuda and M. Stampanoni. High-Resolution Phase-Contrast Imaging of Submicron Particles in Unstained Lung Tissue. AIP Conf. Proc. 1365:384–387, 2011.

28. Seeger W., D. Walmrath, F. Grimminger, S. Rosseau, H. Schütte, H.-J. Krämer, L. Ermert and L. Kiss. Adult respiratory distress syndrome: Model systems using isolated perfused rabbit lungs. Methods Enzymol. 233:549–584, 1994.

29. Soodt T., F. Schröder, M. Klaas, T. van Overbrüggen and W. Schröder. Experimental investigation of the transitional bronchial velocity distribution using stereo scanning PIV. Exp. Fluids 52:709–718, 2012.

30. van der Poll T. and S. M. Opal. Pathogenesis, treatment, and prevention of pneumococcal pneumonia. The Lancet 374:1543–1556, 2009.

31. Wang Y., J. Li, A. Leavey, C. O’Neil, H. Babcock and P. Biswas. Comparative Study on the Size Distributions, Respiratory Deposition, and Transport of Particles Generated from Commonly Used Medical Nebulizers. J. Aerosol Med. Pulm. Drug Deliv. 29:1–9, 2016.

32. Weibel E. R. Morphometry of the Human Lung. Springer-Verlag Berlin Heidelberg, 1963.

33. World Health Organisation. WHO publishes list of bacteria for which new antibiotics are urgently needed. News Release 27 February 2017 (16.11.2018).

34. Xu J., S. L. Murphy, K. D. Kochanek, B. Bastian and E. Arias. Deaths: Final Data for 2016. Natl. Vital Stat. Rep. 67:2018.

35. Yamada Y., A. Koizumi and J. Inaba. A New Method of Casting Human Respiratory Tract for Aerosol Deposition Studies. Radiat. Prot. Dosimet. 79:269–272, 1998.

36. Yanamala N., M. T. Farcas, M. K. Hatfield, E. R. Kisin, V. E. Kagan, C. L. Geraci and A. A. Shvedova. In Vivo Evaluation of the Pulmonary Toxicity of Cellulose Nanocrystals: A Renewable and Sustainable Nanomaterial of the Future. ACS Sustainable Chemistry & Engineering 2:1691–1698, 2014.

37. Zhang Z., C. Kleinstreuer and C. S. Kim. Gas-solid two-phase flow in a triple bifurcation lung airway model. Int. J. Multiphase Flow 28:1021–1046, 2002.

